# Carbapenem-induced β-lactamase-isoform expression trends in *Acinetobacter baumannii*

**DOI:** 10.1101/2024.05.30.596663

**Authors:** Bogdan M. Benin, Trae Hillyer, Noah Aguirre, Yuk Yin Sham, Belinda Willard, Woo Shik Shin

## Abstract

Carbapenem-resistant *Acinetobacter baumannii* (CRAb) is an urgent bacterial threat to public health, with only a few treatment options and a >50% fatality rate. Although several resistance mechanisms are understood, the appearance of these mutations is generally considered stochastic. Recent reports have, however, begun to challenge this assumption. Here, we demonstrate that independent samples of Ab, exposed to different carbapenems with escalating concentrations, show concentration- and carbapenem-dependent trends in β-lactamase-isoform expression. This result, based on the isoforms identified through label-free-quantification LC-MS/MS measurements of cell-free, gel-separated β-lactamases, suggests that the appearance of antibiotic resistance may be somewhat non-stochastic. Specifically, several minor AmpC/ADC β-lactamase-isoforms were found to exhibit both dose- and carbapenem-dependent expression, suggesting the possibility of non-stochastic mutations. Additionally, these also have high sequence similarity to major expressed isoforms, indicating a potential path over which resistance occurred in independent samples. Antibiotic resistance maybe somewhat antibiotic-directed by a hitherto unknown mechanism and further investigation may lead to new strategies for mitigating antibiotic resistance.

**Teaser:** The emergence of antibiotic-resistant β-lactamase proteins from mutations may exhibit patterns based on specific antibiotics.

## Introduction

Carbapenem-resistant *Acinetobacter baumannii* (CRAb) is an urgent threat to public health.(*1*) The CDC chose this classification due to the high mortality rate of these infections, which is in large part due to the current lack of treatment options other than polymyxins, once carbapenems are no longer effective.(*2-6*)

To aid researchers in the development of new treatment strategies and therapeutics, a deeper understanding of the resistance mechanisms utilized by CRAb, and how Ab becomes CRAb, is required. Previously, several reports have suggested that the production of β-lactamases, enzymes that hydrolyze β-lactam antibiotics, is one of the primary mechanisms of resistance.(*6-10*) These enzymes belong to one of four classes (A-D), based on their amino acid sequence and substrate scope.(*11, 12*) More specifically, class A, C, and D β-lactamases contain a catalytic serine residue in their active site and can hydrolyze the majority of β-lactam antibiotics.(*13*) Considering the genome of Ab (ATCC 19606), a common laboratory strain of Ab, genes encoding for class C (*bla*_AmpC_, *bla*_ADC_) and class D (*bla*_OXA_) are present, resulting in the rapid generation of resistance upon β-lactam exposure. Although penicillin binding protein modifications and drug transport proteins may also play a role in the development of CRAb,(*14-16*) several recent reports have suggested that β-lactamases may still be the most significant resistance method.(*6-10*) These studies are generally supported by the observed upregulation of β-lactamases in CRAb whole-proteome studies.(*17-20*) Therefore, understanding the process by which some β-lactamases become carbapenemases would significantly aid researchers in the development of drugs that are less likely to promote the growth of such highly resistant phenotypes.(*21-25*)

To understand the rapid of evolution of drug resistance in bacteria, it is generally considered that either gene transfer or random mutagenesis result in a mutant with higher fitness that survives antibiotic exposure and proliferates to create a more resistant population.(*16*) However, recent studies have shown that this process may not be entirely random as the intrinsic electric field gradients in β-lactamases may be one possible driver for the expression and proliferation of altered enzymes.(*26, 27*) These gradients can correlate with enzymatic activity, providing a spectroscopic handle for resistance prediction. Although the final result of both approaches is the same (*i*.*e*., a resistant population), the initial stages of resistance development are critically different in that one may be subject to predictive modeling and can therefore be influenced.

In an effort to determine the possibility of substrate influenced and non-stochastic β-lactamase expression and evolution, we previously studied the impact of β-lactam exposure (penicillins, cephalosporins, and carbapenems) on the expression of β-lactamases through the use of label-free quantification with LC-MS/MS.(*28*) There, we observed that each β-lactam antibiotic resulted in a unique β-lactamase-isoform resistance profile; however, we were unable to determine if these unique patterns were random or if they were potentiated by the specific type of β-lactam. These observations, overall, fit to the findings of others that suggest that antibiotic resistance may be non-stochastic.

Here, we present a follow-up quantitative proteomic study that seeks to provide additional information to understand the potential non-stochastic nature of β-lactamase mutations and isoform expression patterns. The β-lactamase-isoform expression by Ab19606 in response to exposure to four distinct carbapenems (imipenem, meropenem, doripenem, and biapenem) at three different concentrations was determined through label-free quantification LC-MS/MS, and possible carbapenem- and concentration-dependent trends in β-lactamase expression are identified and reported.

## Results

Starting with a single colony of Ab19606, a total of thirteen flasks with nutrient broth were prepared (**Fig. 1**). Each flask (2L) had a different carbapenem and a concentration of either 1 μM, 5 μM, or 10 μM. These flasks were incubated at 37 °C while stirring at 120 rpm until visible growth was observed (OD units: 0.8∼1.2). A cell-free supernatant (CFS) was then obtained through several rounds of centrifugation and syringe filtering. The β-lactamase proteins within the CFS were then purified and separated utilizing SDS-PAGE gel electrophoresis. Sections belonging to each sample were cut and digested prior to label-free quantification (LFQ) via LC-MS/MS (**Supplementary Table S1, Supplementary Note S1**).

**Figure 1.**
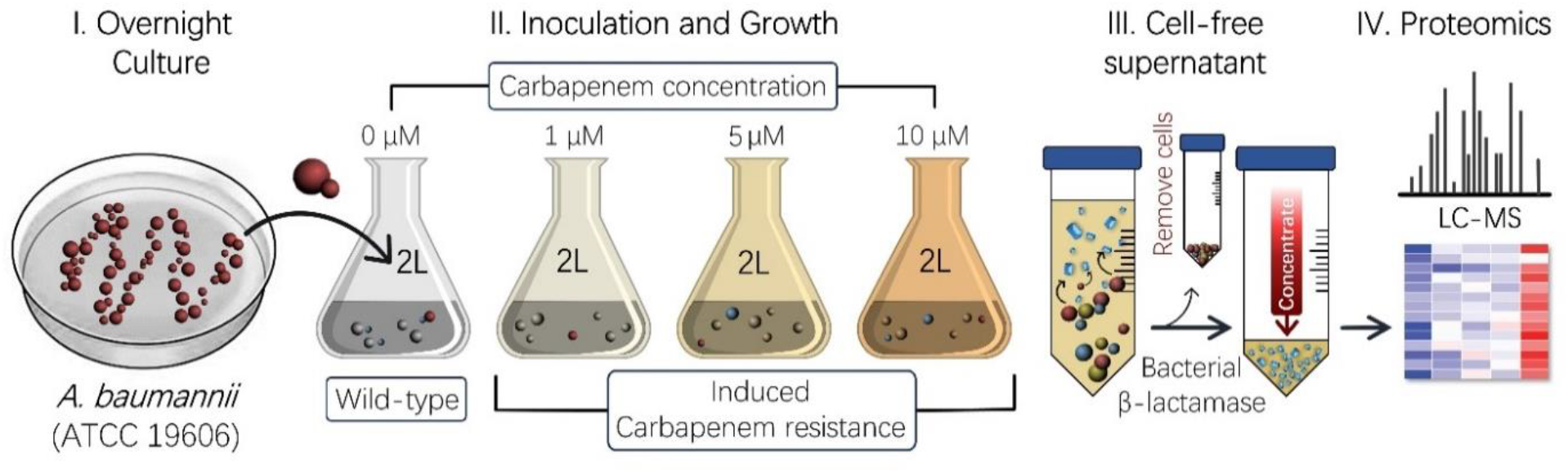
Experimental design for the investigation of carbapenem-dependent and dose-dependent β-lactamase expression patterns.

To study the effect of carbapenem exposure on β-lactamase expression, four distinct carbapenems that are utilized either in the US or abroad were chosen (**Fig. 2a**). These include imipenem, meropenem, biapenem, and doripenem. Importantly, all four share the same core structure (carbapenem core) but have different thio-ether-linked R_2_ sidechains. Effectively, this will result in different 3D structural conformations in stock solutions.

**Figure 2.**
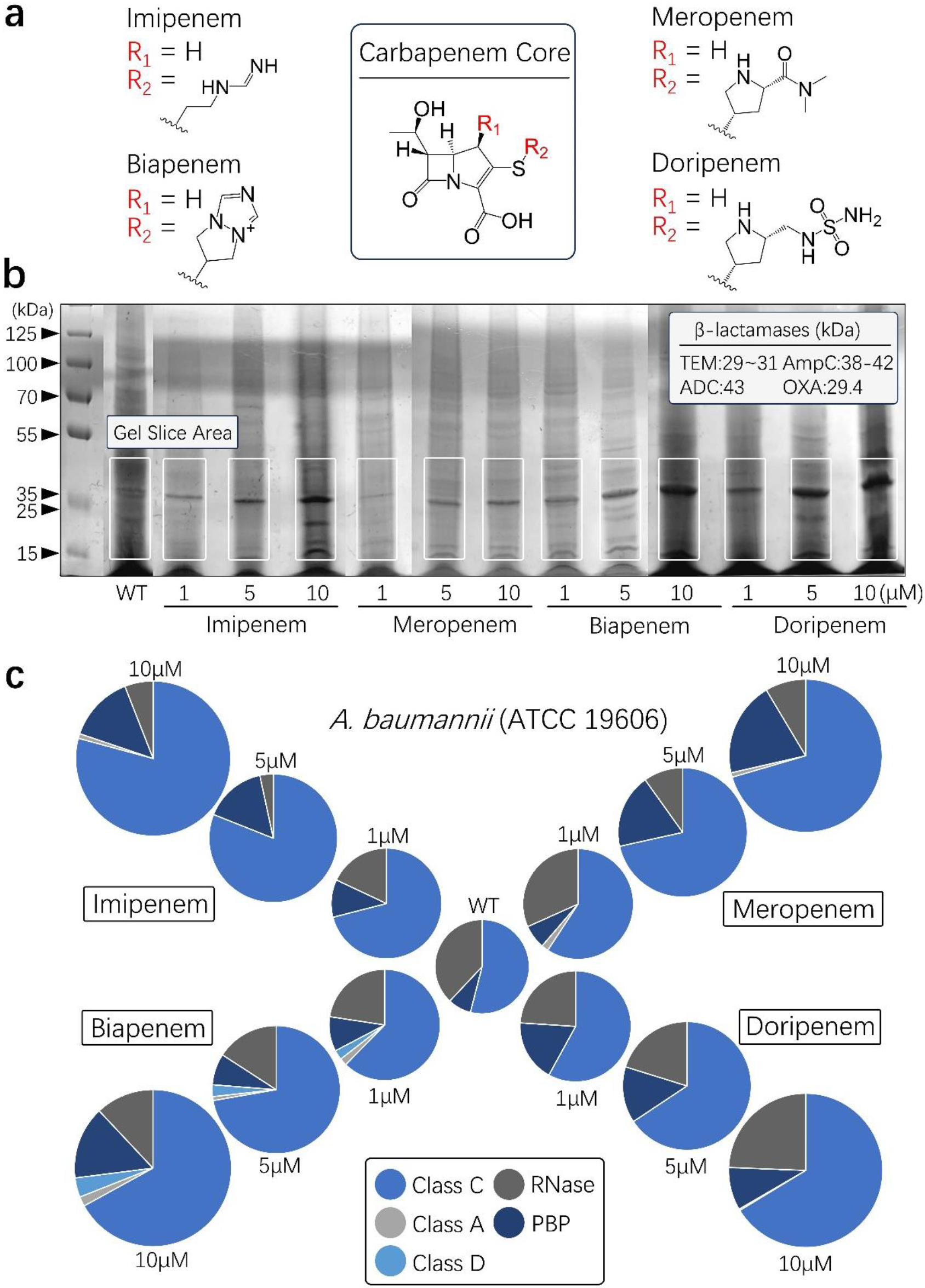
Different β-lactamase profiles are observed depending on which carbapenem has been utilized and at which concentration. **(A)** Four different carbapenems are drawn, highlighting their common core structure and their side-chain variation. **(B)** SDS-PAGE and Coomassie blue staining of cell-free supernatant samples. The possible area of gel containing all classes of β-lactamase proteins (white square) was cut for further proteomics analysis. **(C)** Pie-charts showing the relative expression of different protein classes for each carbapenem-concentration combination.

Additionally, meropenem, biapenem, and doripenem contain methyl groups at the R_1_ position, while imipenem contains hydrogen.

Proteins were identified in MSFragger based on unique peptide recognition. The relative expression of each class of SDS gel-identified proteins is shown in **Figure 2b**. These proteins are categorized into five classes: Class A β-lactamase, Class C β-lactamase, Class D β-lactamase, Zn-containing RNAases, and penicillin binding protein (PBP). Overall, proteins belonging to each class were identified, however, the relative amounts were different for each treatment condition, resulting in unique proteomic profiles.

The unique profiles were further analyzed by only considering the β-lactamase isoforms and their relative expression rates (**Fig. 3**). The majority (>90%) of confirmed β-lactamases were found to belong to class C, with a relatively smaller proportion observed in class D (OXA). Class D, interestingly, demonstrated some signs of carbapenem-dose-dependent expression from Ab exposed to meropenem and imipenem (**Fig. 3b**). However, to further understand if there were any broader effects, and with class C (ADC and AmpC) predominating, a closer look into the identified class C isoforms was carried out (**Fig. 3b, Supplementary Note S1, Supplementary Table S1**).

**Figure 3.**
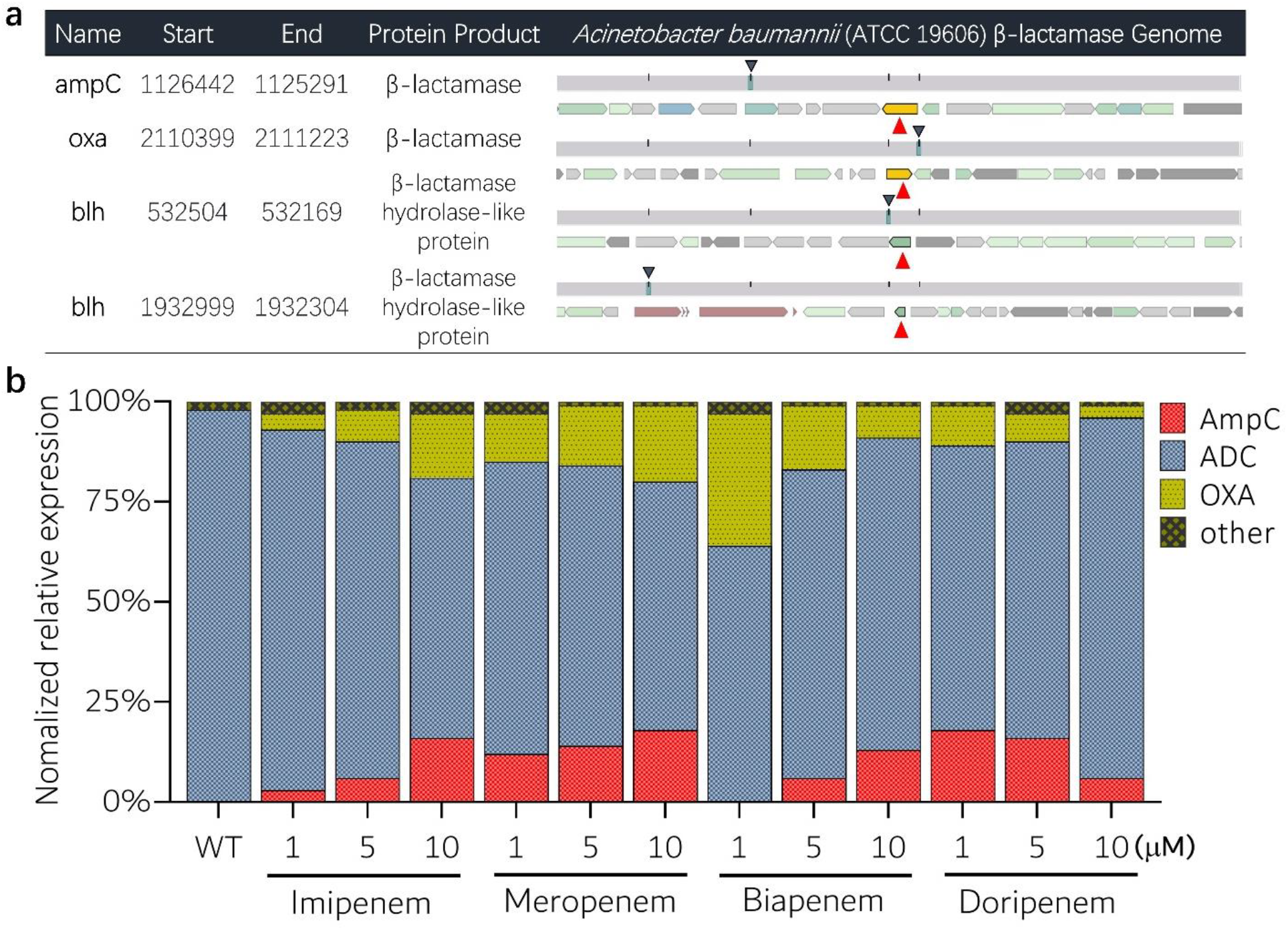
Different β-lactamase expression profiles are observed in independent Ab samples, following treatment with various carbapenems concentrations. **(A)** Ab (ATCC 19606) genomic browser *bla* gene information. **(B)** The relative contribution and relative proportions of each type of identified β-lactamase-isoform to overall β-lactamase expression after carbapenem exposure (1 μM, 5 μM, 10 μM). ADC and AmpC are type C β-lactamases; OXA is type D.

From this, trends in the expression of each uniquely identified Class C isoform based on treatment were investigated, starting with the isoforms that each represented nearly 20% or more of all β-lactamase expression: Q0VTS5, A0A5C0PFX8, A0A858S429 (**Fig. 4a,b**).

**Fig. 4.**
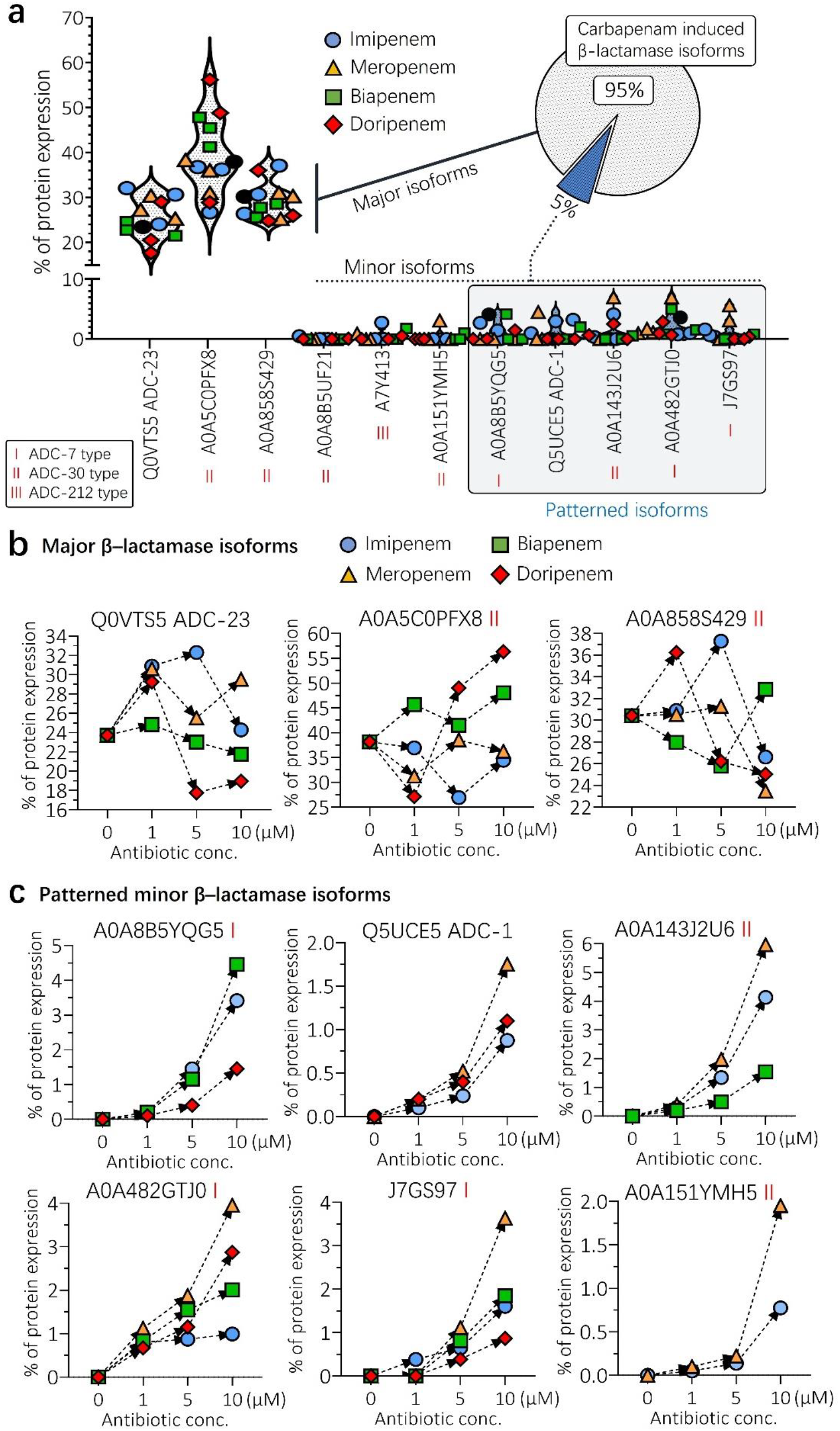
Concentration-dependent trends are observed among non-major class C β-lactamase isoforms. **(A)** All identified isoforms; **(B)** all major isoforms (>95% expression relative to total Class C expression); **(C)** All minor isoforms. Blue circles, green squares, orange triangles, and red diamonds indicate a that the isoform was identified in a sample treated with imipenem, biapenem, meropenem, or doripenem, respectively.

Generally, compared to the control group (Ab19606 wild type), no significant monotonic trend was observed in the expression of these major isoforms (Q0VTS5, A0A5C0PFX8, A0A858S429) (**Fig. 4b**). This suggests that their expression changes do not exhibit a strong correlation with differences in the concentration of the applied carbapenems. Following this, we investigated the potential expression trends among minor isoforms, which generally account for less than 5% of total protein expression (**Fig. 4c**). Surprisingly, the majority of identified minor isoforms exhibited a monotonic expression trend with at least one carbapenem, with five of them showing a pronounced increase in protein expression in response to increased overall carbapenem concentrations. The treatment with imipenem was most frequently associated with a dose-dependent expression trend of β-lactamase protein isoforms, while treatments with doripenem, meropenem, and biapenem also revealed dose-dependent increases in expression albeit for fewer isoforms.

## Discussion

Through our proteomic approach, proteins identified in the Ab ATCC 19606 strain were classified into five major categories (Class A, C, D β-lactamases, RNAases, PBP; **Fig 2**), revealing that Class C β-lactamases were the most abundantly expressed. The other major categories, RNAases and PBP, were not considered further since these may be recognized as an artifact of the experimental design. For example, the concentration dependent increase in PBP is likely indicative of the greater killing of bacteria by carbapenems at higher concentrations. Since PBPs are retained within the cell membrane and not secreted into the extracellular environment like β-lactamases, the increase in PBP identification may result from the release of PBPs into the extracellular space following bacterial death, rather than upregulation. However, this alternative explanation cannot be entirely excluded. We consider the same arguments to be possible explanations for the observation of RNAases in our samples. Instead, by focusing only on the β-lactamase expression, the relative amounts of various β-lactamase classes could be investigated (**Fig. 3**).

The putative Class A β-lactamase (Accession number: A0A1V2V229) was identified, and some concentration dependency was demonstrated. However, since the genome of Ab 19606 does not contain genes encoding for such proteins, this protein may be the result of misidentification. It is, at the current time, difficult to analyze this protein further as all similar records in Uniprot are proteins that have been predicted or inferred from homology and may be subject to change. At the same time, this identified protein is minor and therefore likely plays a limited role in any observed carbapenem resistance.

Surprisingly, the Class D β-lactamases (e.g., OXA) encoded in the Ab 19606 genome were minor contributors, and their expression was found to decrease upon exposure to biapenem in a dose-dependent manner. The reason why the Class D Oxa beta-lactamase, known to be significantly associated with carbapenemase activity, is hardly or not observed in the Ab ATCC 196060 strain has not been elucidated. Although these proteins are similar in molecular weight to both Class C and A proteins, samples were prepared with a minimum amount of polyacrylamide. It is, therefore, possible that a section in which OXA was present, but only in minute amounts, and therefore invisible to the naked eye, may have been missed; however, they would have remained minor contributors to the overall β-lactamase production in the ATCC 196060 Ab strain.

As the primary contributors to identified β-lactamases belonged to Class C, a closer look at possible trends was taken. Here, the major isoforms (those representing 20% or more of Class C) did not exhibit any clear concentration dependence (**Fig. 4b**) although they could be identified in all examined strains (WT and mutant) of Ab ATCC 19606. Instead, pronounced expression trends were mainly observed in minor isoforms (**Fig. 4c**). We observed a tendency for meropenem to induce the greatest relative increase in these minor isoforms as its concentration increases, while imipenem was most frequently associated with a dependent expression trend. Conversely, treatments with doripenem and biapenem resulted in isoform expression rate increases that were either equivalent or relatively lower in correlation.

At this time, none of the physical properties that are readily obtainable from PubChem could be correlated with the observed trends (**Supplementary Fig. S1**). Rather than search for a single needle-in-the-haystack property that could be correlated to our observations, we sought to understand the connection between the various observed protein isoforms. To do so, we consider that these minor isoforms are most likely mutations of the observed major counterparts (Q0VTS5 ADC-23, A0A5C0PFX8, A0A858S429). This was investigated by examining the sequences of each identified isoform and comparing their similarities to each other (**Fig. 5**).

**Fig. 5.**
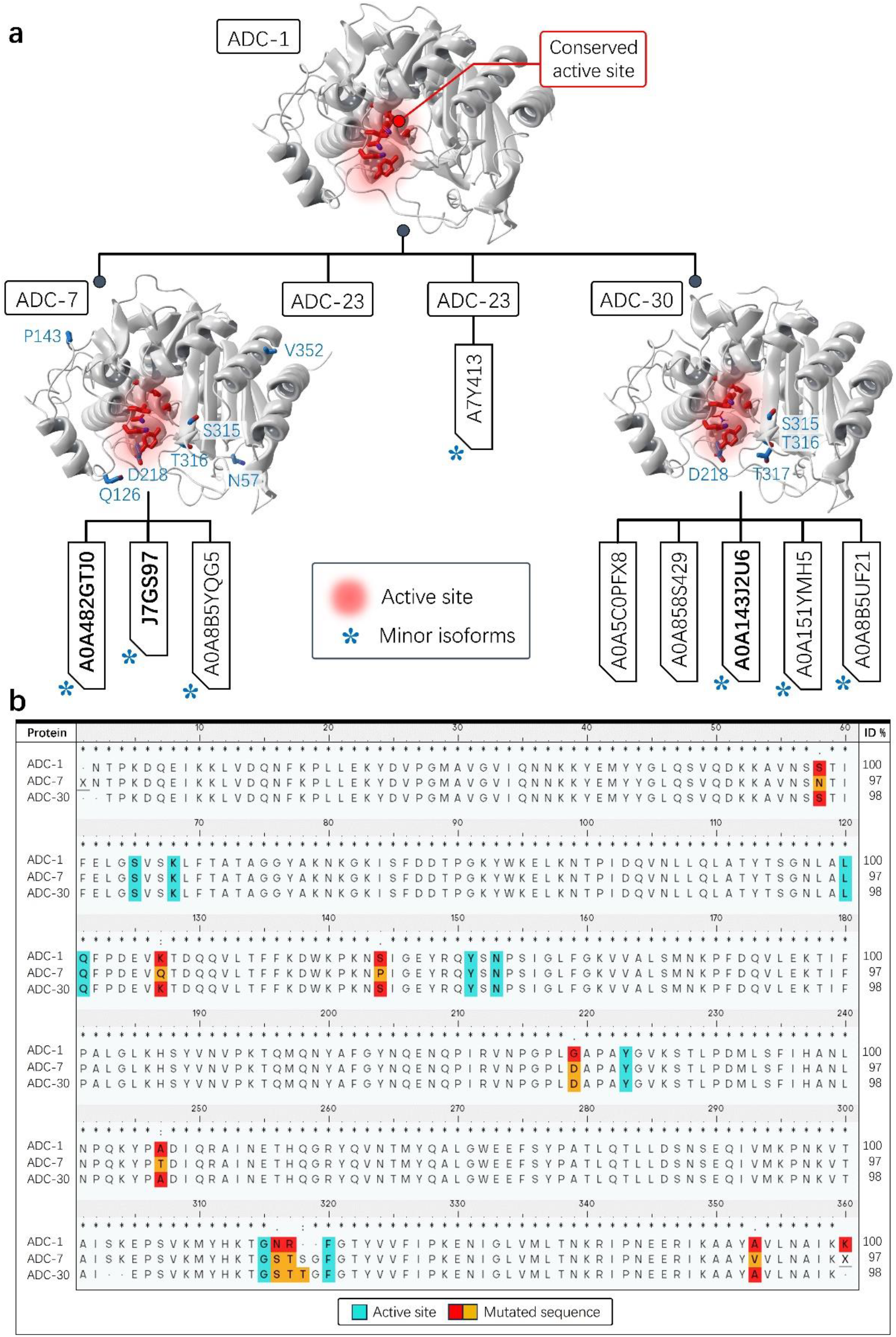
High sequence similarity between identified isoforms demonstrates which mutations have occurred. (**A)** Active binding site and structural comparison of class C β-lactamase ADC-1(PDB ID: 4NET), ADC-7(PDB ID: 4U0T), and ADC-30 (PDB ID: 4U0X). *indicate minor isoforms representing 5% or less of expression. Bolded text represents isoforms that were identified as having a concentration-dependent expression pattern with one or more antibiotics. The active site for ADC isoforms is highly conserved and shown in red. Mutated residues have been indicated in blue. (**B)** Multiple sequence comparison of ADC-1, ADC-7 and ADC-30 β-lactamases. Turquoise indicates residues constituting the active binding site that aligns with the reference sequence (ADC-1). Red and orange represent residues that have mutated to different sequences compared to the reference sequence.

From this, it supports the idea that several of the minor isoforms that follow monotonic trends, are mutants of the major, identified isoforms (Q0VTS5 ADC-23, A0A5C0PFX8, A0A858S429). For example, Q5UCE5 (ADC-1), which was found to increase with imipenem concentration (**Fig. 4c**), has 99% similarity to A0A55C0PXF8, suggesting the two are related by the mutation of several amino acids as well as the possibility that imipenem is in some way potentiating this mutation. Another example of this can be observed with J7GS97 and A0A482GTJ0 (**Fig. 4c**), which both increase in response to all four treatments and are highly similar to ADC-7. However, ADC-7 was not identified by LC-MS/MS in any sample, suggesting that our lab strain may have already had mutations associated with this protein or that numerous mutations took place or that these isoforms are instead associated with the highly similar ADC-30 (**Fig. 5b**).

Again, both sets of observations may suggest that there is a previously unknown, off-site interaction between carbapenems and other regulatory, transcriptional, or translational mechanisms promoting specific mutation patterns. In some cases, such as A0A143J2U6 (**Fig. 4c**), which is a distant mutant from all other Class C isoforms that were identified, trends are observed with all carbapenems utilized in this study. This again seems to point towards a pattern in mutation and expression that cannot be explained by the previous model of completely random mutation resulting in greater fitness and thus increased antibiotic resistance. Instead, we consider that there is the possibility that some mutations are more likely than others due to unknown off-target interactions between antibiotics and bacteria that may increase the likelihood of some phenotypes over others. Our current model of antibiotic resistance is driven, in large part, by random mutation, whereas our proteomic analysis of independent bacterial samples exposed to various carbapenems at three different concentrations demonstrates that some β-lactamase isoforms may be expressed in a dose-dependent manner, calling into question the stochasticity of such mutations. This may suggest that our current model needs revision to include the possibility of off-target interactions or other potentiating factors that favor some mutations over others. Future studies will therefore focus on identifying possible features or properties of these carbapenems that may be associated or correlated with our observed differences in Class C isoform expression as these may also help us determine which drug features may promote specific mutation patterns. This concept, if elucidated, may be of significant benefit to those in drug development as it would offer a new rational tool with which the risk of developing certain phenotypes may be determined.

## Materials and Methods

### Culture Conditions for *A. baumannii*

Nutrient broth was inoculated with *Acinetobacter baumannii* (ATCC 19606) and grown overnight at 37 °C with shaking at 120 rpm. The following day, an aliquot of the bacterial suspension was diluted to an optical density of 0.08-0.1 (600 nm) and spread onto nutrient agar plates and incubated, again, overnight at 37 °C. Two flasks containing 1 L (total 2L) of nutrient broth alone (control) or nutrient broth with varying concentrations of carbapenem antibiotic (1 μM, 5 μM, 10 μM) were inoculated with a single colony. These flasks were then incubated at 37 °C with 120 rpm shaking until cloudiness was observed.

### Supernatant Collection and Purification

From each 1L flask, several 50mL centrifuged tubes were filled and centrifuged twice (8000 x g, 10 min). The entire supernatant was kept and then concentrated using Millipore Sigma Ultra-15 centrifugal filter units with 10 kDa cutoff (Catalog No. UFC901008).

### SDS Gel Electrophoresis

Samples of the concentrated supernatant (7.5 µL) were combined with 2xLaemmli buffer stain (2.5 µL, Bio-Rad) in 1.5 mL microcentrifuge tubes. The samples were heated in a water bath for 10 minutes at 100 °C and then centrifuged (12,000 rpm, 10 min). Samples were cooled on ice prior to gel-loading and then separated at 130V for 2 h.

### Proteomics Analysis

Protein bands were first cut and divided into smaller pieces in order to minimize excess polyacrylamide. These pieces were then washed with water, dehydrated in acetonitrile, reduced with dithiothreitol, and alkylated with iodoacetamide prior to digestion. Digestion was performed in-gel using trypsin (5 μL, 10 μg/mL) in ammonium bicarbonate buffer (50 mM) followed by overnight incubation at room temperature. The resulting peptides were extracted with acetonitrile (30μL, 50%) and formic acid (30μL, 5%), combined, and concentrated in a Speedvac. These samples were finally resuspended in acetic acid (1%) for a final volume of ∼30 μL.

A Bruker TimsTof Pro2 Q-Tof mass spectrometry system operating in positive ion mode, coupled with a CaptiveSpray ion source (both from Bruker Daltonik GmbH, Bremen) was utilized for LC-MS/MS. Samples (1 μL) were injected into the HPLC system, fitted with a 15 cm reversed-phase capillary chromatography column (75 μm inner diameter, packed with C18 ReproSil AQ, 1.9 μm). The peptides were eluted by acetonitrile/0.1% formic acid gradient at a flow rate of 0.3 μL/min and were introduced into the source of the mass spectrometer on-line.

A Parallel Accumulation–Serial Fragmentation DDA method was used to select precursor ions for fragmentation with a TIMS-MS scan (0.60-1.6 Vs/cm^2^ and 100–1,700 *m/z*; ramp time of 166 ms) followed by 10 PASEF MS/MS scans (total cycle time 1.2 s). The MS/MS experiments were performed with collision energies of 20 eV (0.6 Vs/cm^2^) to 59 eV (1.6 Vs/cm^2^). The data was analyzed using all, collected, CID spectra to search a Acinetobacter baumannii (database compiled via UniProt) using MSFragger. Search parameters included the following: precursor mass accuracy of 20 ppm, fragment mass accuracy of 0.05 Da, fully tryptic peptides with 2 missed cleavages allowed, static post-translation modification of carbamidomethylation, and variable post-translational modification of oxidized methionine and protein N-terminal acetylation. Identification was validated to 1% false discovery rate using a decoy database strategy.

### Computational modeling and multiple-sequence analysis

Modeling and structural binding site comparison studies were carried out with the Maestro Schrodinger 2022-2 software package. Computational studies were conducted based on X-ray structures (PDB ID: 4NET, 4U0T and 4U0X). Proteins were prepared through the standard workflow, which includes adding missing sidechains and hydrogens as well as energy minimization using the OPLS-2005 force field.

Our multiple sequence alignment method was used for database search in a straightforward manner. The multiple sequence viewer and alignment tools in Schrodinger package ver. 2022-2 based on classic Smith-Waterman algorithm were used. The comparing sequence data base were provided by UniProt and NCBI Protein Data Bank.

## Supporting information

BioRxiv-SI

## Acknowledgments

Research support for Woo Shik Shin was provided by the Northeast Ohio Medical University, College of Pharmacy. The Northeast Ohio Medical University Department of Pharmaceutical Sciences provided all the necessary research resources.

## Author contributions

T.H. and N.A. grew Ab samples and prepared samples for proteomics. B.W. performed proteomics measurements and peptide identification. B.M.B analyzed the results. B.M.B. and W.S.S. wrote the manuscript. W.S.S. supervised the work. W.S.S., Y.Y.S and B.M.B. designed the project. T.H. and B.M.B. contributed equally to this work. All authors discussed the results and commented on the manuscript.

## Data and materials availability

All analyzed data is available in the main text or the supplementary material and will also be made available from the corresponding author upon reasonable request. The mass spectrometry proteomics data have been deposited to the ProteomeXchange Consortium via the PRIDE partner repository with the dataset identifier.

