## Supplementary material for "Carbapenem-induced β-lactamase-isoform expression trends in *Acinetobacter baumannii*": BioRxiv-SI: BioRxiv-SI_Shin.docx

**for**

**Table S1. β-lactamase protein identification by unique peptide sequences.**

| Accession ID | Gene ID | Unique Peptides |
| --- | --- | --- |
| A0A858S429 | HIN86_0549 | 2 |
| Q0VTS5 | ADC-23 | 2 |
| A0A5C0PFX8 | blaADC | 1 |
| A0A482GTJ0 | blaADC | 1 |
| Q5UCE5 | blaAmpC | 2 |
| A0A8B5YQG5 | blaADC | 1 |
| A0A143J2U6 | blaAmpC | 1 |
| J7GS97 | N/A | 3 |

**Supplementary Note S1. Identified β-lactamase isoform amino acid sequences.** Unique, identified sequences are red.

A0A1V2V229_9GAMM

Protein MW: **45274**

Amino Acid Composition: **A32 C2 D22 E23 F17 G22 H4 I25 K38 L50 M10 N25 P22 Q18 R7 S38 T24 V28 W4 Y4**

MKNFLTKKMY ASRALFFAVG VMALNVSPIV ASTAHAAAEQ KMDNLSTKLS SIFADKPIEA SLFSPQFLGQ VSITQIQKIV DDLKASLGTL KSINVSNGSG TIDFEKGELP VSISLDEQGQ INTLWFSAPH FKTVSLDEMV KGLHQNAVGK TSLLVVVDNK PVVVENDKTP MAVGSTFKLL VLKAYEDAIK KGELKRETIV SLKEKNRSLP TGVLQNLPDD TPVNLELLAQ LMIQISDNTA TDSLIDELKK PRIEALSPRN IPMLTTRELF QLIDPSNEKL KNKFKTGTKS ARLEALSDLD KLPLPSVSDI GKSATWQDAE WYMSANEICP LLESAQDAPA LNSSLNPLFK NLNWQKIGFK GGSEYGVINF SVIGKTQKGH KVCAVFTANG NEPQPESKLA ILFTGILQAV DSMNP

Q0VTS5_adc-23

Protein MW: **43155**

Amino Acid Composition: **A23 C1 D15 E17 F19 G23 H3 I23 K39 L33 M9 N28 P25 Q24 R6 S22 T24 V25 W3 Y21**

MRFKKISCLL LPPLFIFSTS IYAGNTPKEQ EVKKLVDQNF KPLLDKYDVP GMAVGVIQNN KKYEIYYGLQ SVQDKKAVNS STIFELGSVS KLFTATAGGY AKAKGKISFD DTPGKYWKEL KNTPIDQVNL LQLATYTSGN LALQFPDEVQ TDQQVLTFFK DWKTKNAIGE YRQYSNPSIG LFGKIVALSM NKPFDQVLEK TIFPPLHLKN SYVNVPKTQM QNYAYGYNQE NQPIRVNPGP LDAPAYGVKS TLPDMLTFIN ANLNPQKYPK DIQRAINETH QGFYQVGTMY QALGWEEFSY PASLQTLLDS NSEQIVMKPN KVTAISKEPS VKMFHKTGST NGFGTYVVFI PKENIGLVML TNKRIPNEER IKAAYAVLNA IKK

*A0A143J2U6_ampC*

Protein MW: **43114**

Amino Acid Composition: **A22 C1 D17 E16 F20 G24 H3 I24 K40 L34 M9 N25 P25 Q22 R7 S24 T24 V23 W3 Y20**

MRFKKISCLL LSPLFIFSTS IYAGNTPKDR EIKKLVDQNF KPLLDKYDVP GMAVGIIQNN KKYETYYGLQ SVQDKKAVSS STIFELGSVS KLFTATAGGY AKTKGTISFD DAPGKYWKEL KNTPIDQVNL LQLATYTSGN LALQFPDEVK TDQQVLTFFK DWKPKNSIGE YRQYSNPSIG LFGKVVALSM NKPFDQLLEK TIFPDLGLKH SYVNVPKTQM QNYAFGYNQE NQPIRVIPGP LDAPAYGVKS TLPDMLKFIN ANLNPQKYPK DIQRAINETH QGFYQVGTMY QALGWEEFSY PAPLQTLLDS NSEQIVMKPN KVTAISKEPS VKMFHKTGST NGFGTYVVFI PKENIGLVML TNKRIPNEER IKAAYAVLNA IKK

*A7Y413_ampC*

Protein MW: **43326**

Amino Acid Composition: **A23 C1 D16 E17 F18 G23 H4 I22 K38 L34 M10 N28 P23 Q23 R8 S24 T23 V25 W3 Y21**

MRFKKISCLL LSPLFIFSTS IYAGNTPKDQ EIKKLVDQNF KPLLEKYDVP GMAVGVIQNN RKYEMYYGLQ SVQDKKAVNS STIFELGSVS KLFTATAGGY AKNKGKISFD DTPGKYWKEL KNTPIDQVNL LQLATYTSGN LALQFPDEVK TDQQVLTFFK DWKPKNSIGE YRQYSNPSIG LFGKVVALSM NKPFDQVLEK TIFPALGLKH SYVNVPKTQM QNYDFGYNQE NQPIRVNPGP LDALAAYGVK STLPDMLSFI HANLNPQKYP ADIQRAINET HQGRYQVNTM YQALGWEEFS YPATLQTLLD SNSEQIVMKP NKVTAISKEP SVKMYHKTGS TNGFGTYVVF IPKENIGLVM LTNKRIPNEE RIKAAYAVLN AIKK

*A0A151YMH5_AWW73*

Protein MW: **43118**

Amino Acid Composition: **A21 C1 D13 E18 F20 G24 H3 I22 K36 L36 M9 N28 P23 Q26 R6 S25 T24 V25 W3 Y20**

MRFQKISCLL LSPLFIFSTS IYAGNTLKEQ EIKKLVDQNF KPLLEKYNVP GMAVGIVQNN KKYEIYYGLQ SVQDKKAVNS STIFELGSVS KLFTATAGGY AKTKGTLSFE DTPGKYWKEL KNTPIDQVNL LQLATYTSGN LALQFPDEVQ TDQQVLTFFK DWKPKNSIGK YRQYSNPSIG LFGKVVALSM NQSFDQVLEK TIFPDLGLKH SYVNVPKLQM QNYAFGYNQE NQPIRVNPGP LDAPAYGVKS TLPDMLSFIN ANLNPQKYPA DIQRTINETH QGFYQVGTMY QALGWEEFSY PAPLQTLLDS NSEQIVMKPN KVTAISKEPS VKMFHKTGST NGFGTYVVFI PKENIGLVML TNKRIPNEER IKAAYAVLNA IKK

Q5UCE5_bla(ampC)

Protein MW: **43236**

Amino Acid Composition: **A23 C1 D14 E17 F18 G23 H4 I22 K38 L33 M10 N28 P24 Q23 R9 S24 T23 V25 W3 Y21**

MRFKKISCLL LSPLFIFSTS IYAGNTPKDQ EIKKLVDQNF KPLLEKYDVP GMAVGVIQNN KRYEMYYGLQ SVQDKKAVNS STIFELGSVS KLFTATAGGY AKNKGKISFD DTPGKYWKEL KNTPIDQVNL LQLATYTSGN LALQFPDEVK TDQQVLTFFK DWKPKNSIGE YRQYSNPSIG LFGKVVALSM NKPFDQVLEK TIFPALGLKH SYVNVPKTQM QNYAFGYNQE NQPIRVNPGP LGAPAYGVKS TLPDMLSFIH ANLNPQKYPA DIQRAINETH QGRYQVNTMY QALGWEEFSY PATLQTLLDS NSEQIVMKPN KVTAISKEPS VKMYHKTGST NRFGTYVVFI PKENIGLVML TNKRIPNEER IKAAYAVLNA IKK

A0A8B5UF21_blaADC

Protein MW: **43215**

Amino Acid Composition: **A23 C1 D15 E17 F18 G22 H4 I22 K38 L33 M10 N27 P23 Q25 R7 S25 T24 V25 W3 Y21**

MRFKKISCLL LSPLFIFSTS IYAGNTPKDQ EIKKLVDQNF KPLLEKYDVP GMAVGVIQNN KKYEMYYGLQ SVQDKKAVNS STIFELGSVS KLFTATAGGY AKNKGKISFD DTPGKYWKEL KNTSIDQVNL LQLATYTSGN LALQFPDEVQ TDQQVLTFFK DWQPKNPIGE YRQYSNPSIG LFGKVVALSM NKPFDQVLEK TIFPALGLKH SYVNVAKTQM QNYAFGYNQE NQPIRVNPGP LDAPAYSVKS TLPDMLKFIH ANLNPQKYPT DIQRAINETH QGRYQVNTMY QALGWEEFSY PATLQTLLDS NSEQIVMKPN KVTAISKEPS VKMYHKTGST SGFGTYVVFI PKENIGLVML TNKRIPNEER IKAAYAVLNA IKK

A0A482GTJ0_blaADC

Protein MW: **43126**

Amino Acid Composition: **A23 C1 D16 E16 F19 G23 H4 I22 K39 L33 M10 N26 P25 Q23 R6 S23 T24 V26 W3 Y21**

MRFKKISCLL LSPLFIFSTS IYAGNTPKDQ EIKKLVDQNF KPLLEKYDVP GMAVGVIQNN KKYEMYYGLQ SVQDKKAVNS STIFELGSVS KLFTATAGGY AKNKGKISFD DTPGKYWKVL KNTPIDQVNL LQLATYTSGN LALKFPDEVQ TDQQVLTFFK DWKPKNPIGE YRQYSNPSIG LFGKVVALSM NKPFDQVLEK TIFPALGLKH SYVNVPKTQM QNYAFGYNQE NQPIRVNPGP LDAPAYGVKS TLPDMLSFIH ANLNPQKYPA DIQRAINETH QGFYQVNTMY QALGWEEFSY PATLQTLLDS NSEQIVMKPN KVTAISKEPS VKMYHKTGST TGFGTYVVFI PKENIGLVML TNKRIPNEER IKAAYAVLDA IKK

*A0A5C0PFX8_blaADC*

Protein MW: **43168**

Amino Acid Composition: **A23 C1 D15 E17 F18 G23 H4 I22 K39 L33 M10 N27 P24 Q23 R7 S23 T25 V25 W3 Y21**

MRFKKISCLL LSPLFIFSTS IYAGNTPKDQ EIKKLVDQNF KPLLEKYDVP GMAVGVIQNN KKYEMYYGLQ SVQDKKAVNS STIFELGSVS KLFTATAGGY AKNKGKISFD DTPGKYWKEL KNTPIDQVNL LQLATYTSGN LALQFPDEVK TDQQVLTFFK DWKPKNSIGE YRQYSNPSIG LFGKVVALSM NKPFDQVLEK TIFPALGLKH SYVNVPKTQM QNYAFGYNQE NQPIRVNPGP LDAPAYGVKS TLPDMLSFIH ANLNPQKYPA DIQRAINETH QGRYQVNTMY QALGWEEFSY PATLQTLLDS NSEQIVMKPN KVTAISKEPS VKMYHKTGTT TGFGTYVVFI PKENIGLVML TNKRIPNEER IKAAYAVLNA IKK

A0A858S429_HIN86_05495

Protein MW: **43179**

Amino Acid Composition: **A22 C1 D15 E17 F18 G23 H4 I22 K38 L33 M10 N28 P25 Q24 R7 S25 T22 V25 W3 Y21**

MRFKKISCLL LSPLFIFSTS IYAGNTPKDQ EIKKLVDQNF KPLLEKYDVP GMAVGVIQNN KKYEMYYGLQ SVQDKKAVNS STIFELGSVS KLFTATAGGY AKNKGKISFD DTPGKYWKEL KNTPIDQVNL LQLATYTSGN LALQFPDEVQ TDQQVLTFFK DWKPKNPIGE YRQYSNPSIG LFGKVVSLSM NKPFDQVLEK TIFPALGLKH SYVNVPKTQM QNYAFGYNQE NQPIRVNPGP LDAPAYGVKS TLPDMLSFIH ANLNPQKYPA DIQRAINETH QGRYQVNTMY QALGWEEFSY PATLQTLLDS NSEQIVMKPN KVSAISKEPS VKMYHKTGST NGFGTYVVFI PKENIGLVML TNKRIPNEER IKAAYAVLNA IKK

D0CDC9_ACIB2

Protein MW: **31989**

Amino Acid Composition: **A15 C8 D11 E16 F13 G18 H19 I17 K21 L36 M5 N11 P11 Q21 R11 S7 T11 V12 W4 Y9**

MQKTKIMIHK IHHLKCGSMC PVCAPLFGQR GWKAEIVCHC LLVETDRGLV LIDTGFGLQD YLHMQQRLGS LVKRLGKIEP NLEFSAIQQI QKLGFHPKDV QHIFVTHLDF DHAGGISDFP HATVHVLATE YNAAQLPNFK GKLRYRTNQY KQHRYWNFVE YQQGEKWFNL EKVKGLPLFQ DEILMVPLLG HSAGHCGIAI KQQNQWLLFC GDAYYSHLEL NPANKLRSLT LLEKTFAEDN EKRLINLKKL QHLAQHEPTI EIICAHDPHE LRRYQT

D0CG83_ACIB0

Protein MW: **35369**

Amino Acid Composition: **A31 C2 D15 E13 F16 G23 H14 I16 K28 L29 M3 N19 P16 Q18 R4 S17 T16 V23 W3 Y14**

MKKLFVALGL IMGSLHISYA EPASAQQVPG YYHHQFGNYR ITSLLDGTIY LDPKLFKNLS PAEKTKILTK YAAVNEKGIQ TSVNAFLVDD GKSLTLVDSG AASCFGPQLG SIAKNLELAG YQLANVKTVL LTHLHPDHAC GIAQNGKAVF PNATIYAHER EADYWLNPEN EKTVPADKKE NYLGTVKNVK AALAPYQAKK AFKTFKDGDV IQGFEVINTQ GHTPGHHSFR LKSKGQQIVF VGDIVHSHSL QFDAPKTGVD FDVNSEQAIN TRLKMFAEVS NKQQWVAAPH LPFPGIGHVY KVNAEQYQWI PLYFNNSLDK

A0A836YKZ8_J568_0973

Protein MW: **34148**

Amino Acid Composition: **A29 C2 D15 E12 F15 G22 H13 I15 K27 L25 M2 N19 P15 Q19 R4 S18 T16 V24 W3 Y14**

MGGLHVSYAE PASAQQVPGY YKHQFGNYRI TSLLDGTIYL DPKLFKNLSQ AEKTKILTKY AAVNEKGVQT SVNAFLVDDG KSLTLVDSGA SSCFGPQLGS IAKNLELAGY QLANVKTVLL THLHPDHVCG IAQNGKAVFP NATIYAHERE ADYWLNPASE KTVPADKKEN YLGTVKNIKA ALAPYQAKKA FKTFKDGDVI QGFEVINTQG HTPGHHSFRL KSKNQQIVFV GDIVHSHSLQ FDAPKTGVDF DVNSEQAINT RLKMFAEISN KQQWVAAPHL PFPGIGHVYK VNAEQYQWIP LYFNNSVDK

A0A013RR17_J658_3771

Protein MW: **31744**

Amino Acid Composition: **A26 D13 E16 F11 G16 H4 I19 K30 L27 M7 N8 P15 Q17 R2 S21 T19 V23 W4 Y10**

MKFFKLTASV LATATALTVT SSAFAQDLKI QSFLAKPEHF GVTSTLIEGD KEVLLVNAQF SKSEALRIAA DILDSGKTLK TIFVSYGDPD YYFGLDVFKQ YFPNVQIIAT PETVKHIQDT QALKVKYWGP QMGANAPSKI IMPQAYTAKT LKLENESIEI KGKKELTYLW IPSSKAVVGG IPVSSGIHLW MADTPKVKDR NEVIQTLENI KALQPQTVIP AHMVEGAPQG LDAVNFSIDY LKSYEKAVKV SKNAGELSQL MQKQYPTLKS VDSLELGAKV VKGEMQWP

A0A086HV98_pbpG

Protein MW: **35381**

Amino Acid Composition: **A43 C1 D13 E12 F6 G23 H2 I13 K10 L33 M9 N28 P15 Q8 R19 S36 T30 V22 W3 Y9**

MSILLSLSST SFAELVNNPS SGSTGTASLN WSAADASQLL NDEDDEPTPQ GSTSVTTTLR GSNAPRVITS APKVAPIRDT VGYNAQPSVS ARAALVMDAQ TGEVLYSKNT NASVPIASIT KLMTAVVTAD ARLNMSEEIT LEQIDFAGAG GKNSSSTLRV GDKMNRAEVL LFALMKSENP AAAALARTYP GGRAAFVAAM NAKARALGMN ATHYYESTGL DPRNVSSARD LGILASTASQ YGLIRQFSTT PTYDFNLGYR VLKSNNTNAL VRNGGWNINL SKTGYINEAG RCVVMHTTVN SRPAVIVLLG EPSTQARNND ATNLLGWLSN LPKRI

J7GS97_ACIBA

Protein MW: **43230**

Amino Acid Composition: **A22 C1 D15 E17 F20 G23 H4 I22 K38 L32 M10 N28 P25 Q24 R6 S23 T23 V26 W3 Y21**

MRFKKISCLL LSPLFIFSTS IYAGNTPKDQ EIKKLVDQNF KPLLEKYDVP GMAVGVIQNN KKYEMYYGFQ SVQDKKAVNS STIFELGSVS KLFTATAGGY AKNKGKISFD DTPGKYWKEL KNTPIDQVNL LQLATYTSGN LALQFPDEVQ TDQQVLTFFK DWKPKNPIGE YRQYSNPSIG LFGKVVALSM NKPFDQVLEK TIFPALGLKH SYVNVPKTQM QNYAFGYNQE NQPIRVNPGP LDAPAYGVKS TLPDMLSFIH ANLNPQKYPA DIQRAINETH QGFYQVNTMY QALGWEEFSY PATLQTLLDS NSEQIVMKPN KVTAISKEPS VKMYHKTGST NGFGTYVVFI PKENIGLVML TNKRIPNEER IKAAYVVLNA IKK

J9XPV4_oxa51

Protein MW: **13279**

Amino Acid Composition: **A10 D6 E9 F5 G8 H2 I5 K12 L11 M3 N4 P5 Q8 R4 S4 T7 V8 W3 Y3**

NALIGLEHHK ATTTEVFKWD GKKRLFPEWE KDMTLGEAMK ASAIPVYQDL ARRIGLELMS KEVKRVGYGN ADIGTQVDNF WLVGPLKITP QQEAQFAYKL ANKTLPFSQK VQDEVQS

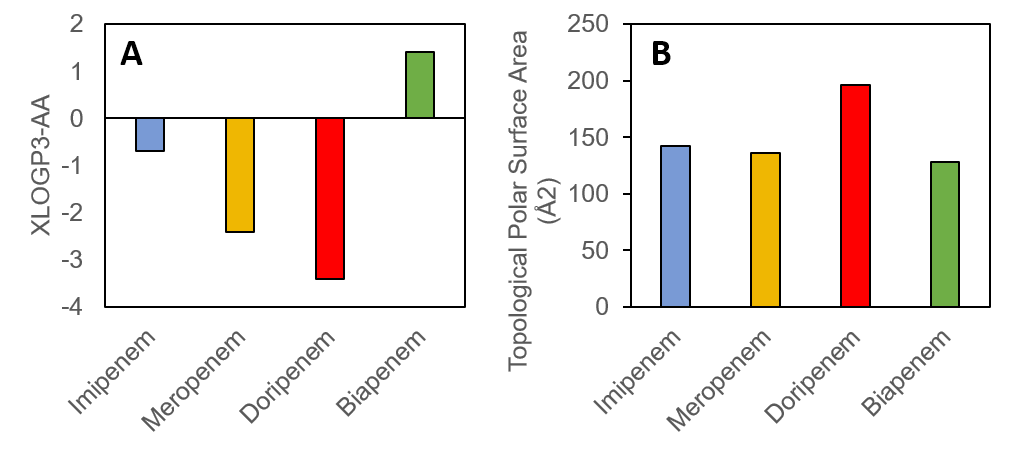

**Fig. S1. Calculated physical properties.** (**A**) Octanol-water partition coefficient. (**B**) Surface area. All values were taken from PubChem.
